# A Phosphoinositide Interacting Protein Coordinates Stress Precursor Activities

**DOI:** 10.1101/2025.08.18.670812

**Authors:** Pooja Roy, Blake A. Rose, Suhita Ray, Meg A. Schaefer, Ventakasai R. Dogiparthi, Suyong Choi, Nicholas T. Woods, Kyle J. Hewitt

## Abstract

Samd14 is crucial for cell signaling and survival in mouse models of acute anemia. Samd14 has an N-terminal actin capping protein (CP) and a C-terminal sterile alpha motif (SAM) to coordinate stem cell factor/Kit and erythropoietin receptor signaling pathways during terminal differentiation of red blood cell precursors. Here we present new findings that Samd14 expression is needed to maintain balanced autophagy in red blood cell precursors following acute anemia. Autophagy gene signatures and protein levels are markedly altered within the context of acute anemia when red blood cell precursors accelerate the process of erythropoiesis. Samd14 interacted via its SAM domain with phosphatidylinositol 3-phosphate (PI3P) which is an integral component of endosomal and autophagic membranes. We tested PI3Ps role in red blood cell differentiation using a small molecule inhibitor of the Class III PI 3-kinase VPS34, which is the sole kinase responsible for PI3P genesis. SAR405 treatment blocked red blood cell formation. In the absence of Samd14, higher doses of VPS34 inhibition were required to block erythroid differentiation. Given the critical roles of autophagy in normal differentiation, Samd14’s stress-dependent activation and roles in autophagy suggest that this mechanism is needed to maintain progenitor levels and balance the production of healthy, mature red blood cells.

**Key Points:**

- Autophagy genes/proteins are deregulated in red blood cell progenitors after hemolysis-induced acute anemia.
- The anemia-activated Samd14 protein interacts with the PI3P lipid moiety which is integral to endosomal and autophagic membrane functions.
- VPS34 inhibition blocks terminal erythroid maturation in a Samd14-dependent manner.

## Introduction

During normal red blood cell differentiation, cell size decreases and organelles are cleared to streamline hemoglobin production. Intracellular components such as mitochondria and ribosomes are recycled by the well-known process of autophagy (Zhang *et al*, 2015). Autophagy-related genes and protein expression are elevated throughout erythroid cell commitment to control cell size and maintain protein quality control (Kang *et al*, 2012; Khandros *et al*, 2012). The autophagy machinery is influenced by cytokine signaling pathways – including stem cell factor (SCF)/Kit (Ertmer *et al*, 2007; Shi *et al*, 2025) – which regulate aspects of cell differentiation including self-renewal and survival (Kroemer *et al*, 2010). Among the important steps, the PI-3-kinase vacuolar protein sorting-34 (VPS34) enzyme produces phosphatidylinositol 3-phosphate (PI3P) where it accumulates to initiate autophagosome membranes (Obara & Ohsumi, 2008). Autophagy mechanisms play a role in red cell disorders including β-thalassemia and myelodysplastic syndrome (Khandros *et al*., 2012; Lechauve *et al*, 2019; Mortensen *et al*, 2010).

While autophagy is required for the terminal stages of normal erythropoiesis, it is also a critical mechanism of cellular responses to stress (Kroemer *et al*., 2010). To wit, how do red blood cell progenitors integrate autophagy mechanisms required for normal erythropoiesis with stress-dependent changes in environmental conditions (e.g. in acute anemia)? The acceleration of erythropoiesis needed to restore homeostasis in anemia is carried out by a unique population of stress erythroid progenitor (SEP) cells (Paulson *et al*, 2020). After lineage commitment by stress signals, SEPs undergo a period of rapid expansion characterized by a requirement for unique signaling cues (Lenox *et al*, 2005; Perry *et al*, 2007). SCF/Kit and downstream activation of ERK and AKT/mTOR are critical mediators of SEP expansion and activity (Knight *et al*, 2014; Ray *et al*, 2020; Subramanian *et al*, 2005). Following treatment with the red blood cell-lysing chemical phenylhydrazine (PHZ), mice bearing loss-of-function alleles of the Kit receptor have severe delay in SEP colony forming activity (Perry *et al*., 2007). Kit and mTOR signaling, along with autophagy, are integrated into processes of erythroid progenitor activation, expansion and differentiation to relieve acute anemia (Knight *et al*., 2014).

Anemia is common, but knowledge about anemia recovery and key mechanisms involved in balancing the distinct requirements of “stress erythropoiesis” are incomplete (Paulson *et al*., 2020). Prior work identified GATA factor-dependent transcriptional activation of genes predicted to control acute responses to anemia (Hewitt *et al*, 2017; Zhou *et al*, 2025; Zhou *et al*, 2023). The sterile alpha motif-containing protein-14 (Samd14) is anemia-regulated via an intronic GATA factor-regulated enhancer (Hewitt *et al*., 2017). Samd14 boosts the erythroid system’s potential for regeneration (Schaefer *et al*, 2023). Samd14 has a conserved sterile alpha motif (SAM) which is required for anemia-dependent SCF/Kit signaling, survival and progenitor colony-forming activity (Ray *et al*., 2020; Ray *et al*, 2022). Sterile alpha motifs (SAMs) provide interfaces for molecular interactions. SAMs often self-interact or can form SAM-SAM oligomers and polymers, and SAMs can bind DNA, RNA and lipids (Ray & Hewitt, 2023). While the SAM domain of Samd14 is required for stress progenitor cell activity and cell signaling downstream of Kit activation, heterologous proteins interacting with this domain or other molecular interactors are not known.

In this study, we investigated the process of autophagy in stress erythropoiesis and the role of Samd14. We identified a SAM-dependent interaction between Samd14 and the autophagy-associated PI3P phospholipid. We link the Samd14-PI3P molecular association with a need to rebalance deregulated autophagy during expansion and differentiation of stress erythroid progenitors.

## Results

### Molecular and phenotypic attributes of autophagy are Samd14-sensitive in stress erythroid progenitors

Samd14 potentiates higher SEP responses to SCF in acute anemia (Hewitt *et al*., 2017; Hewitt *et al*, 2025; Ray *et al*., 2022), however the mechanism is unclear. Phenylhydrazine (PHZ)-induced red blood cell hemolysis is a reliable *in vivo* model to test cellular and molecular responses to acute anemia, as it rapidly induces anemia and stimulates SEP expansion in mouse spleen. To probe Samd14 functions in acute anemia, we conducted global phospho-proteomics in lineage (Lin)^-^Kit^+^ cells from PHZ treated mice which were acutely stimulated with the Kit ligand SCF for 25 minutes (**Figure 1A**). 3,057 phospho-proteins were detected in both treatment groups (**Figure 1B**, **Table 1**). As expected, SCF stimulation of SEPs increased the phosphorylation of MAPK components (**Figure 1C**). The transcriptional repressor capicua (CIC), a previously-described hyperphosphorylated protein downstream of receptor tyrosine kinase activation (Jimenez *et al*, 2012), was phosphorylated after SCF stimulation (**Figure 1C**). All 4 biological replicates exhibited significant increases in the Y185 phosphorylation of mitogen activated protein kinase 1 (MAPK1, also known as ERK2) (**Figure 1D**). Enriched phospho-proteins post-SCF included proteins with known kinase and phosphatase activities, and components of the translational machinery (**Figure 1E**). Conversely, the top cellular process downregulated by Kit signaling was autophagy (**Figure 1F**), consistent with known roles of this pathway inhibiting autophagy (Ertmer *et al*., 2007; Shi *et al*., 2025). We compared autophagy phospho-proteins from cells isolated from wild type (WT) mice to cells isolated from a Samd14 enhancer knockout genetic background (S14^ΔE/ΔE^) which are unable to recover from severe anemia (Hewitt *et al*., 2017). The decrease in autophagy-associated phospho-proteins following SCF stimulation was significantly less in S14^ΔE/ΔE^ cells (**Figure 1F**).

**Figure 1:**
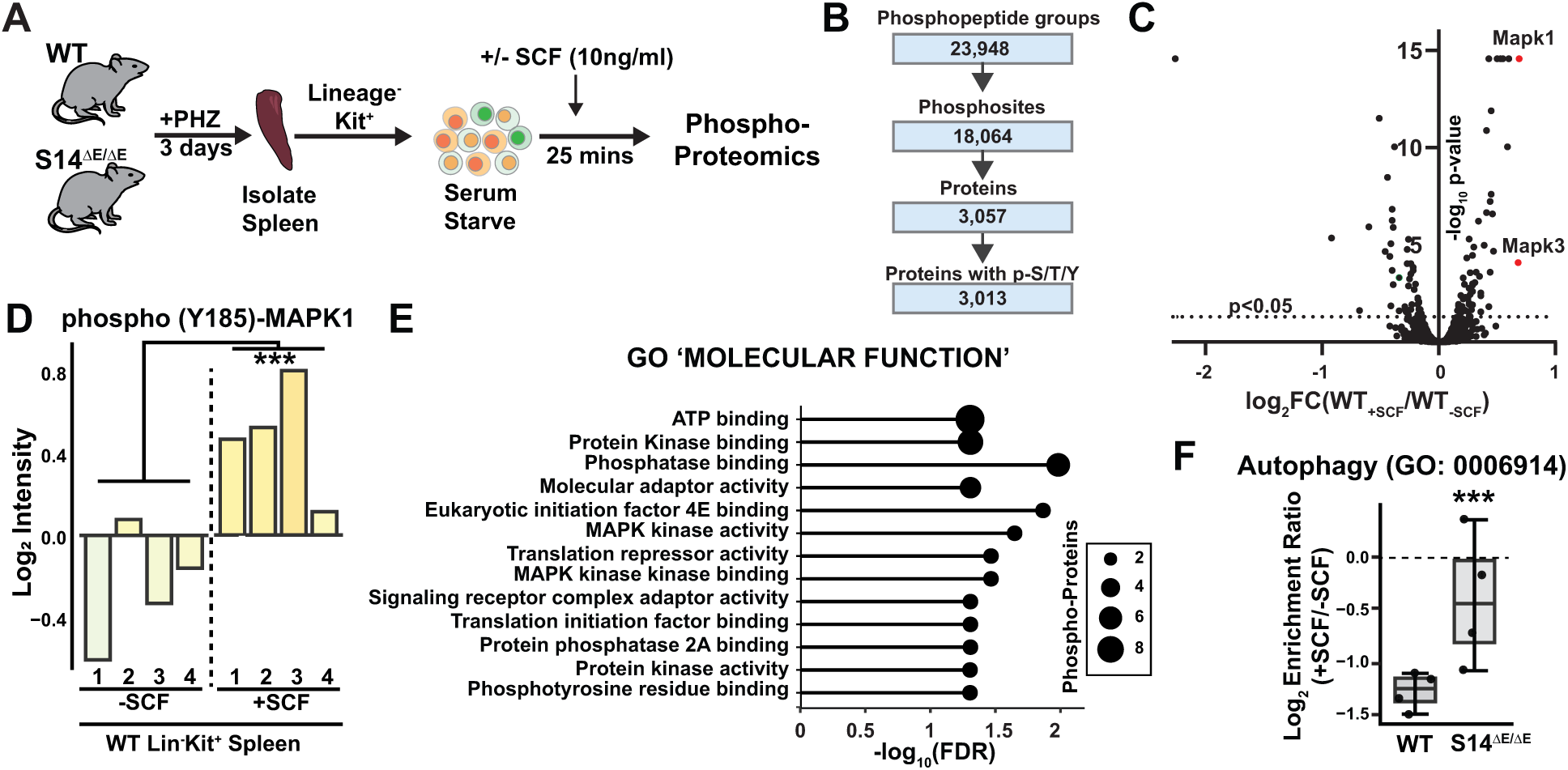
Kit pathway activation inhibits autophagy. A) Experimental layout of primary Kit^+^ cells isolated from phenylhydrazine (PHZ)-treated WT and Samd14^ΔE/ΔE^ mice and SCF stimulated stimulated (10 ng/mL) prior to phosphoproteomic analysis (N=4 mice per genotype). B) Numbers of phospho-proteins and phospho-sites identified. C) Volcano plot representing Log_2_-fold change (FC) of WT (control) vs. WT (+SCF) phospho-proteins. D) Bar graph of Log2-normalized signal intensity of phospho (Y185)-MAPK phospho-peptide in WT control vs. WT SCF-treated cells. E) Gene ontology (GO) analysis of phosphoprotein-associated molecular functions enriched after SCF treatment. F) Box and whisker plot of GO enrichment ratios (-/+SCF) derived from phospho-proteins involved in autophagy which were depleted in SCF-treated samples. Error bars represent standard deviation (SD). ***p<0.001 (two-tailed unpaired Student’s t test).

We next compared global changes to Samd14-sensitive phospho-proteins before and after Kit activation. The total number of phospho-proteins enriched or depleted in S14^ΔE/ΔE^ cells compared to WT cells were 127 (untreated), 133 (SCF-treated) and 234 (combined) (fold change<1.5, P<0.05) (**Figure 2A**, **Table 2**). Phospho-Samd14 was 22-fold higher in untreated WT cells vs S14^ΔE/ΔE^, with phosphorylations at S77, S165, S173, S179 and S267 residues. Phospho-proteins depleted in Samd14-deficient cells were involved in regulation of autophagy (Prkaa1, Rps6kb1, Pdpk1), mTOR signaling (Rps6kb1, Pdpk1, Prr5, Prkaa1), and protein translation (Eif3d, Denr) (**Figure 2B**). Phosphorylation of the regulatory subunit p101 of the Class IB phosphoinositide 3-kinase complex (Pik3r5) - which lacks catalytic activity (Andrews *et al*, 2007) - increased in Samd14-deficient SEPs (**Figure 2C**). To link differentially-phosphorylated peptides to their cognate kinase activate, we conducted a kinase enrichment analysis (KEA3) (Kuleshov *et al*, 2021). Top kinase hits included canonical effectors of Kit signaling (Akt, Mapk and mTOR), and several kinases involved in apoptosis and autophagy (Csnk2a1 and Chek2) not previously linked to Kit signaling (**Figure 2D**). Gene ontology pathway analysis revealed that phosphorylations of Mtor pathway, AMPK signaling and autophagy-related proteins were distinct in Samd14-deficient cells (**Figure 2E**).

**Figure 2:**
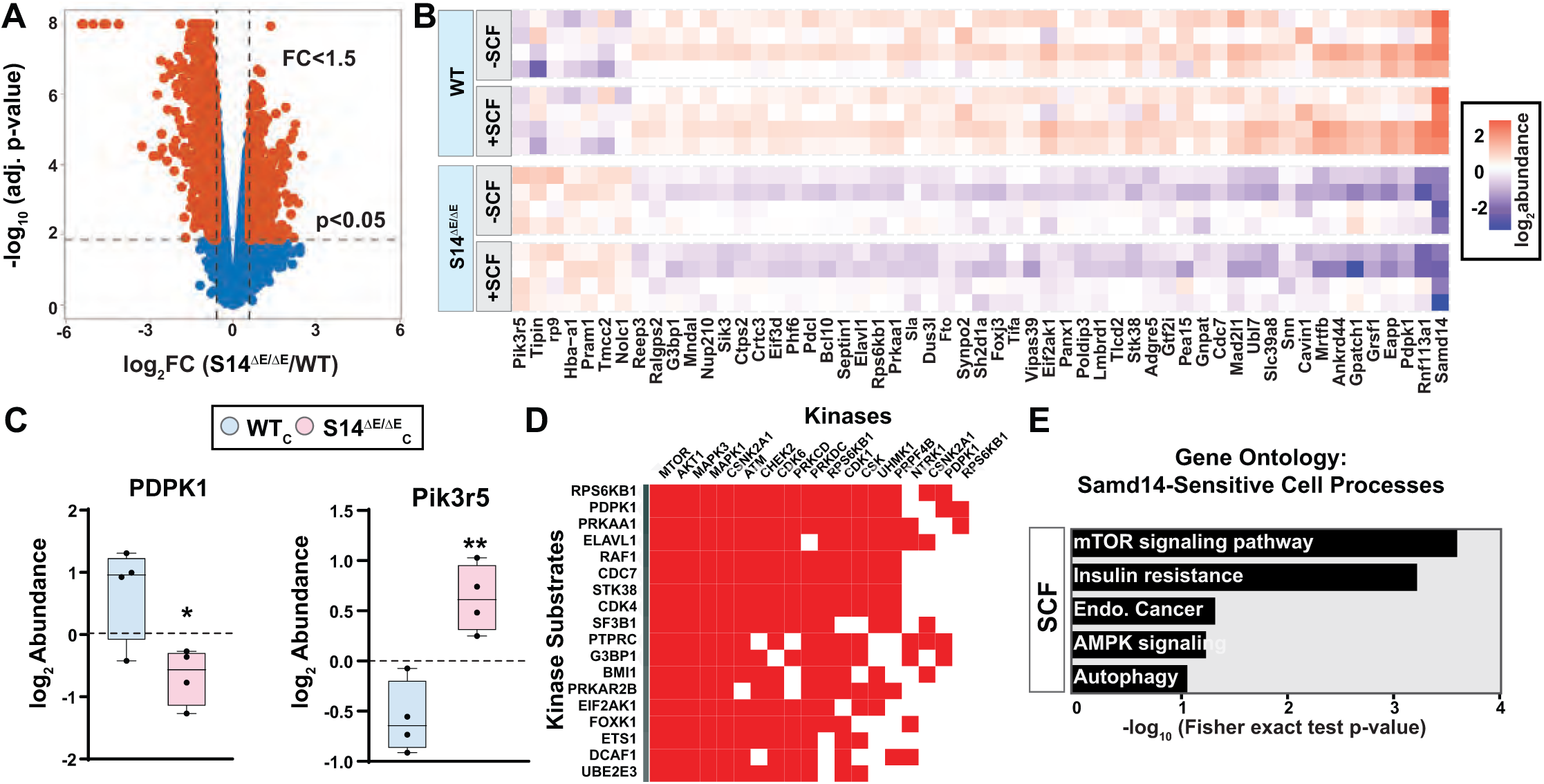
Samd14-sensitive phosphoproteins are involved in stress-dependent autophagy. A) Volcano plot representing Log_2_-fold change of identified phospho-sites between WT and Samd14^ΔEnh/ΔEnh^ cells. 3,193 phospho-sites were significantly up- or down-regulated. FC=fold change >1.5 (N=4 mice per genotype). B) Heat map depicts the top 40 differentially-enriched phosphoproteins in WT vs Sand4^ΔEnh/ΔEnh^ cells in both control (serum-starved) and SCF-treated samples. C) Box and whisker plots of normalized abundances of 3-phosphoinositide-dependent kinase 1 (PDK1) and regulatory subunit p101 of the Class IB phosphoinositide kinase (Pik3r5) in control WT and S14^ΔE/ΔE^ cells (n=4). Error bars represent SD. *p<0.05; **p<0.01 (two-tailed unpaired Student’s t test). D) Clustergram of predicted kinases upstream of SAMD14-regulated phosphoproteins using Kinase Enrichment Analysis version 3 (KEA3) in RokaiXplorer. E) GO of the top 5 Kyoto Encyclopedia of Genes and Genomes (KEGG)-enriched signaling and metabolic pathways elevated upon SCF-treatment in WT vs S14ΔE/ΔE erythroid progenitors. Endo=endometrial cancer.

Since our data suggested Samd14-sensitive regulation of autophagy in acute anemia, we evaluated the expression of key autophagy genes in acute anemia vs. controls. In Lin^-^ CD71^+^ cells, autophagy-regulated transcripts were downregulated in acute anemia (PHZ) vs. controls (PBS), including the transcription factor EB (*Tfeb*) which was 5.2-fold downregulated (**Figure 3A**). While *Map1lc3b* transcripts were down 1.3-fold, the levels of lipidated LC3-II were higher in acute anemia vs. controls (**Figure 3B**). Protein levels of phospho-EIF2α (Ser 51), which downregulates global protein synthesis in response to various stress stimuli and promotes autophagy (B’Chir *et al*, 2013; Talloczy *et al*, 2002) was upregulated in anemia versus steady state (**Figure 3B)**. Protein level of phospho-Ampk (Thr 172), which promotes autophagy and red blood cell homeostasis by inhibiting mTOR signaling, (Foretz *et al*, 2010; Hawley *et al*, 1996), is higher in splenic erythroblasts after anemia (**Figure 3B**). This is accompanied by upregulation of the mTORC1 substrate, phospho-4EBP1 (Thr 37/46) after anemia (**Figure 3B**). These findings illustrate that acute anemia shifts the cellular requirements such that SEPs expansion requires fine-tuning the balance between mTORC1 and autophagy pathway activation.

**Figure 3.**
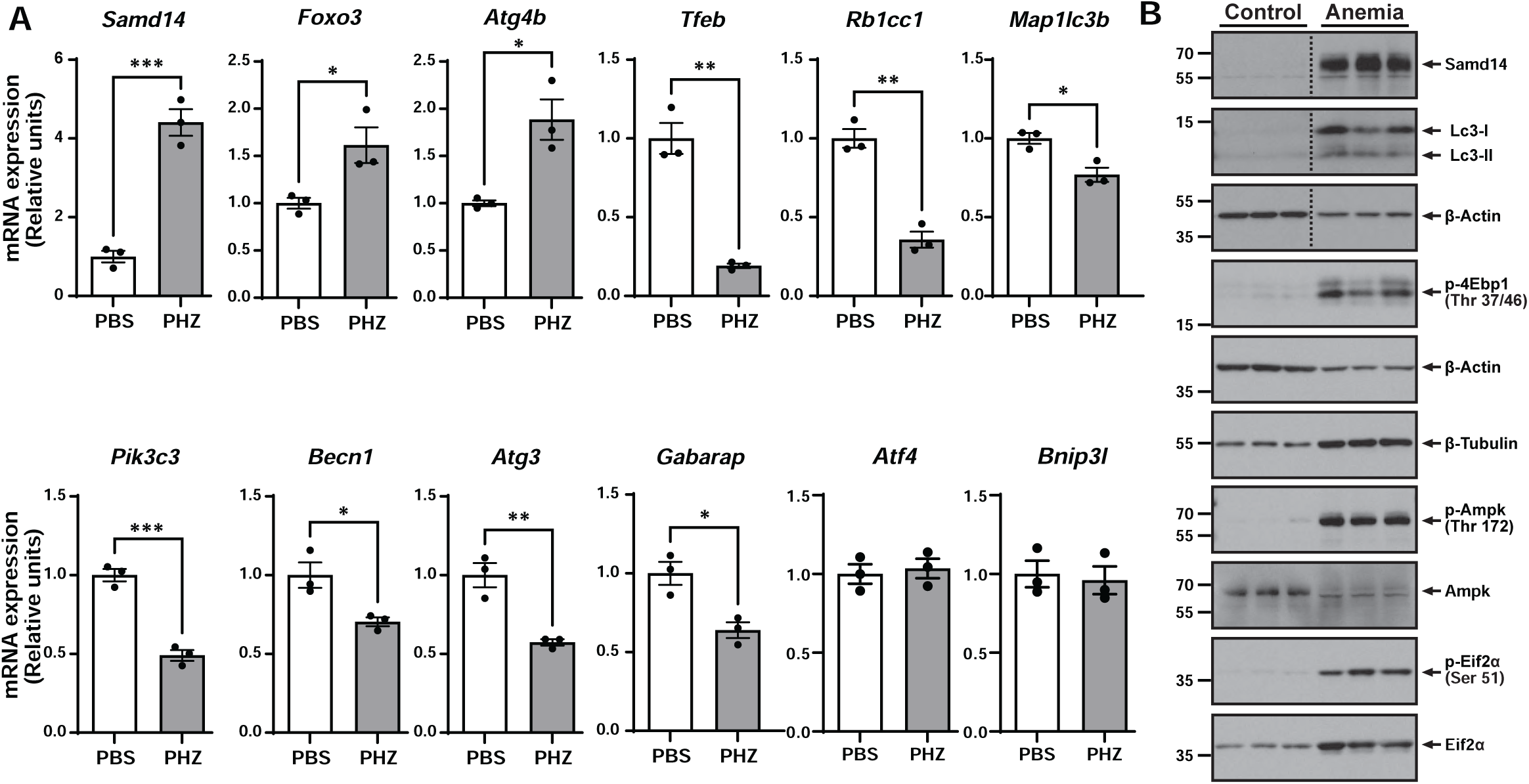
Autophagy-related proteins are deregulated in acute anemia. A) qRT-PCR in murine splenic erythroblast cells (Lineage^-^CD71^+^) 3 days after acute hemolytic anemia. Error bars represent standard error of mean (SEM). *p<0.05; **p<0.01; ***p<0.001 (two-tailed unpaired Student’s t test). (B) Western blotting of Lineage^-^CD71^+^ cells isolated from control (PBS-treated) and anemia (PHZ) spleens for autophagy-related proteins. N=3 mice per experimental group.

### A phosphatidylinositol 3-monophosphate (PI3P) lipid present in autophagosome membranes interacts with the Samd14 SAM

Sterile Alpha Motifs (SAMs) in SAM-containing proteins play widely varying roles in cell biology, including autophagy, but no known SAM-dependent protein-protein interactions of Samd14 have been detected (Ray *et al*., 2022). As approximately 40% of SAMs self-associate (Knight *et al*, 2011), we tested whether Samd14 self-associates via its SAM. Mouse GATA1- and Samd14-null G1E cells were co-infected with retrovirus containing FLAG-tagged and HA-tagged Samd14 (**Figure 4A**). Co-infected cells expressed HA- and FLAG-tagged Samd14 at similar levels (**Figure 4B**). Immunoprecipitation (IP) with anti-HA antibody followed by western blotting with anti-FLAG antibody revealed that FLAG-tagged Samd14 failed to pull down the HA-tagged Samd14 (**Figure 4C**). Instead, abundant FLAG-tagged Samd14 remained in the IP supernatant (**Figure 4D**). This holds true even when lysis conditions varied from 1% NP-40 to 0.1% NP-40 (**Figure 4E**). We conclude that Samd14 does not self-dimerize under conditions we tested.

**Figure 4.**
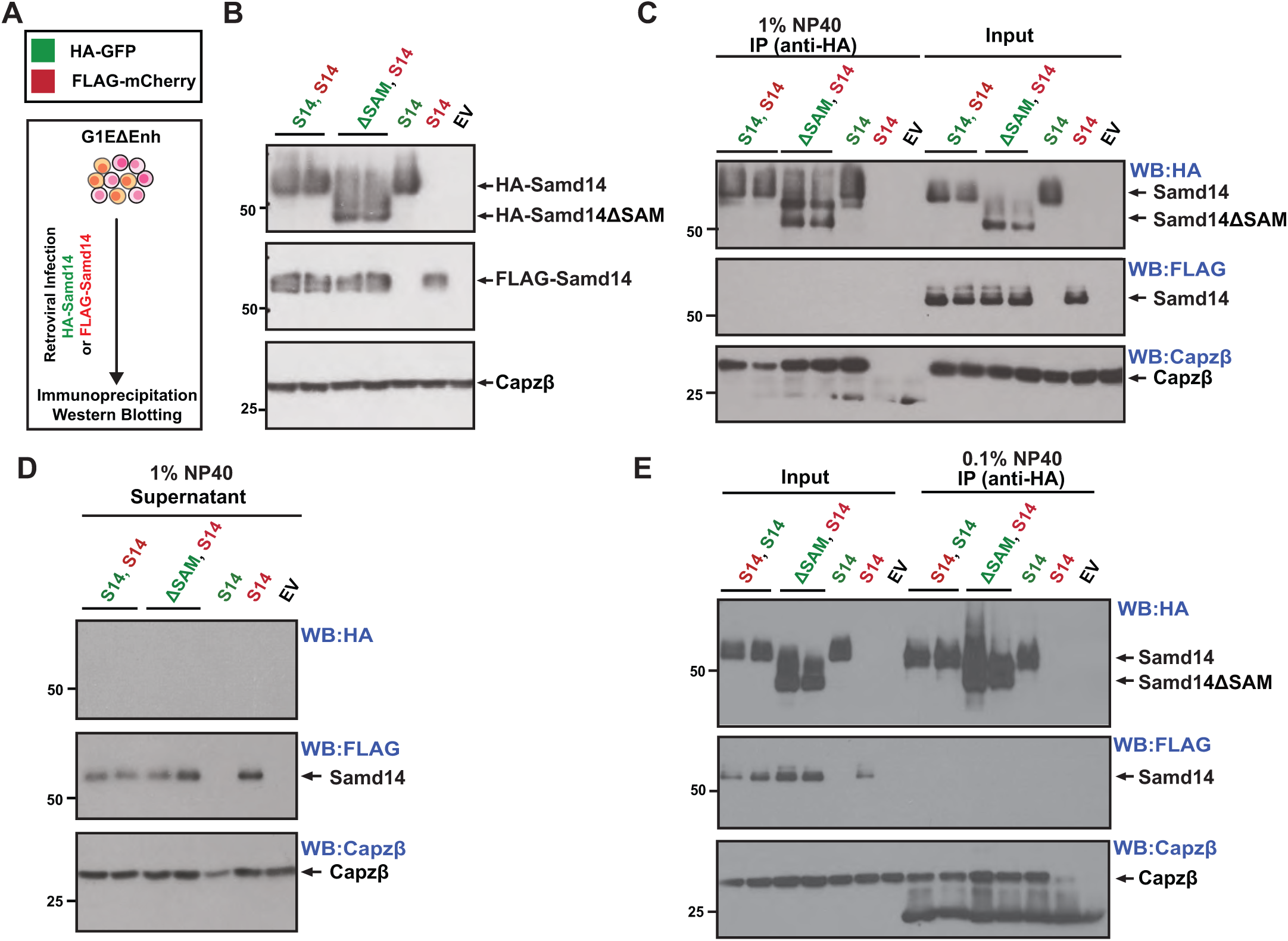
SAM domain of Samd14 does not self-dimerize. A) G1E-ΔEnh cells were co-infected with retrovirus containing FLAG-tagged Samd14 along with HA-tagged Samd14 or HA-tagged SAM domain-deleted mutant of Samd14. Protein self-association was tested by carrying out immunoprecipitation with anti-HA beads followed by western blotting with anti-FLAG antibody. B) Western blotting of total protein lysates from G1E-ΔEnh cells retrovirally infected with expression constructs for HA-Samd14, FLAG-Samd14 and doubly infected cells. C) Western blotting of HA immunoprecipitated protein lysates (lysed in 1% NP-40) from HA-Samd14, FLAG-Samd14, and co-infected cells. D) Western blotting of supernatant fractions after HA immunoprecipitation from HA-Samd14, FLAG-Samd14, and co-infected cells. E) Western blotting of HA immunoprecipitated protein lysates (lysed in 0.1% NP-40) from HA-Samd14, FLAG-Samd14, and co-infected cells.

Phosphoinositides represent potential molecular interactors with the Samd14-SAM (Barrera *et al*, 2003). Sequence alignment of Samd14-SAM from different vertebrate species revealed an evolutionarily conserved polybasic motif (PBM) located at the C-terminal end of its SAM domain (**Figure 5A**). PBMs are clusters of basic amino acids that many proteins utilize to engage in electrostatic interactions with the inositol head group of phosphoinositide (Kaadige & Ayer, 2006; Wurmser & Emr, 2002). Alpha fold predicted a solvent side motif within Samd14 SAMs 5^th^ alpha helix (**Figure 5B**). We hypothesized that a SAM-dependent Samd14 interaction with phosphoinositides facilitates erythroid progenitor cell activity in anemia recovery. To test this, we performed a protein-lipid overlay blot using affinity purified protein lysates from G1E cells retrovirally infected with expression constructs containing HA-tagged full length Samd14, SAM deleted (S14ΔSAM), capping protein binding (CPB) deleted (S14ΔCPB), both domains deleted (S14ΔCS or a point mutation in the putative polybasic motif (S14R381Q) (**Figure 5C**). Full length Samd14 binds PI3P with high specificity, while also binding at lower affinity to PI4P and PI5P (**Figure 5D**). The Samd14-PI3P interaction was undetectable using the SAM-deleted Samd14 or the SAM/CPB-deleted Samd14, however it was maintained using the CPB-only deleted Samd14 (**Figure 5D**). While prior reports revealed a CP-PI3P interaction that controls actin assembly (Mi *et al*, 2015; Schafer *et al*, 1996), our results show that Samd14 interacts independently with PI3P and CP through distinct domains and that CP cannot facilitate Samd14’s interaction with PI3P. The Samd14 PBM-mutant (S14R381Q) protein bound PI3P at much lower affinity (**Figure 5D**). To confirm this interaction was present in cells, we conducted a proximity ligation assay (PLA) after expressing Samd14 (or deletion/mutant constructs) in cells lacking endogenous Samd14 expression (**Figure 5E**). Compared to full length Samd14, the number of PLA puncta decreased by 3.3-fold in the absence of the SAM domain and by 1.9-fold in a PBM mutant (S14R381Q) (**Figure 5F**). To test the importance of the Samd14-PI3P interaction on the function of erythroid progenitor cells, we conducted a colony formation assay (CFU) in lineage-negative cells isolated from fetal liver of Samd14 knockout embryos (E14.5). As previously reported, the SAM domain is required for Samd14’s promotion of progenitor activity (Ray *et al*., 2020). Progenitors expressing the S14R381Q mutant Samd14 formed 1.9-fold fewer burst forming unit-erythroid colonies (p=0.01) compared to cells expressing full length Samd14 (**Figure 5G**). Samd14-SAM binds to the endosome/autophagosome-associated PI3P phosphoinositide via R381 and PI3P binding is needed for Samd14 to promote erythroid colony formation (**Figure 5H**).

**Figure 5.**
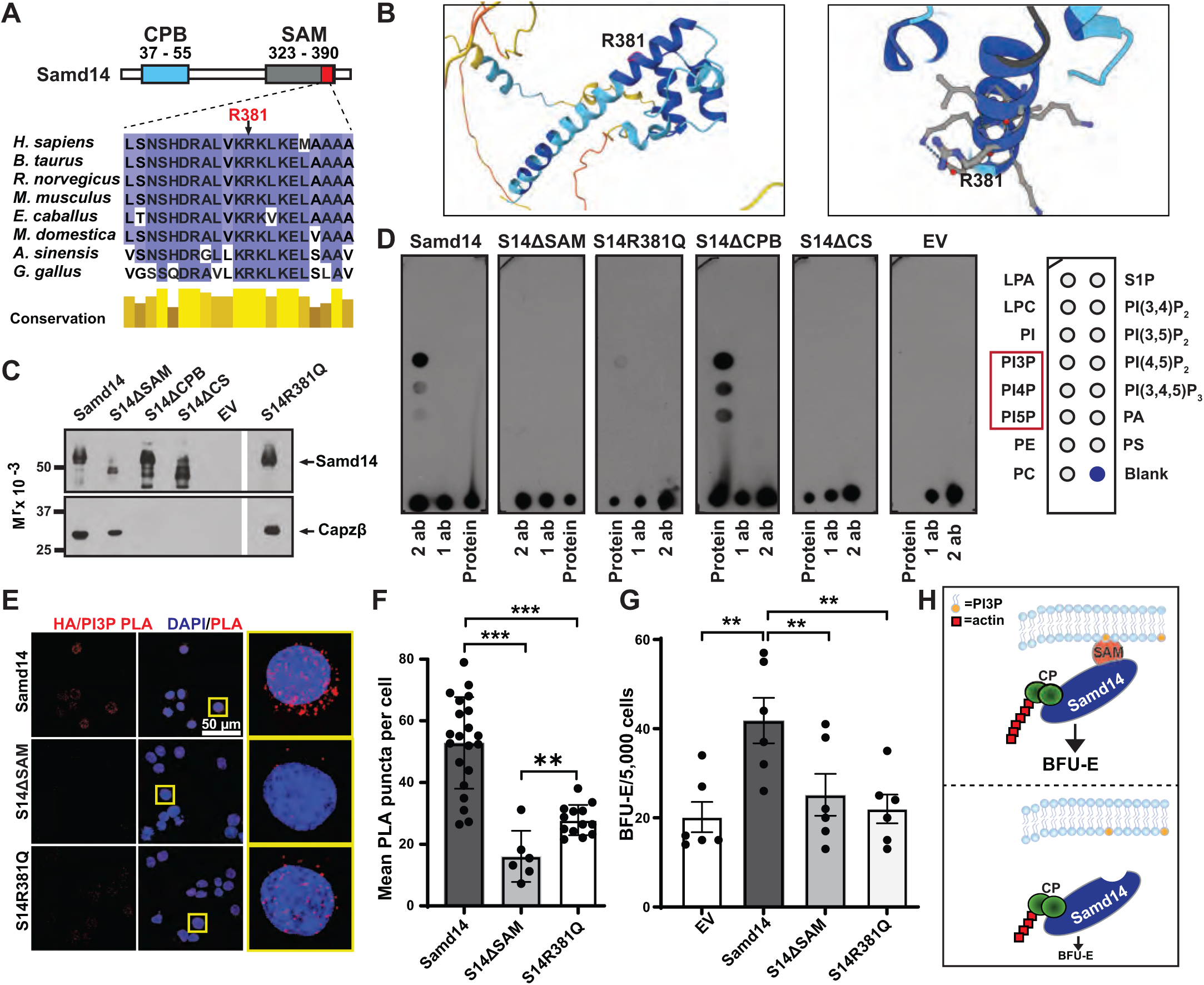
Samd14 interacts with Phosphatidylinositol 3-Phosphate (PI3P) using an arginine residue located within its SAM domain. A) A ‘KRK’ polybasic motif (PBM) in the Samd14-SAM is evolutionarily conserved among vertebrates (visualized using JalView). B) Location of R381 in the 5^th^ SAM alpha helix, predicted by AlphaFold3 (Abramson *et al*, 2024). C) Western blotting of protein lysates from HA affinity purified retrovirally-infected G1E cells, representing protein fractions used for PIP strip assay. D) Images of PIP strips incubated with HA-Samd14 (full length), HA-S14ΔSAM (missing SAM domain), HA-S14ΔCPB (missing CP-interaction domain), HA-S14ΔSAMΔCPB (missing both domains), HA-S14R381Q (PBM mutant), or an empty vector control lysate. The bottom of the strip was dotted with protein, 1° and 2° antibody as positive controls. E) Proximity ligation assay (PLA) in G1E cells expressing HA-Samd14 (full length), HA-S14ΔSAM (missing SAM domain), HA-S14R381Q (PBM mutant) with antibodies against HA and PI3P. Nucleus was stained with DAPI. F) Quantification of average PLA puncta observed per cell in an image field using ImageJ (Andy’s algorithm). Each data point is representative of an image field (50-100 cells were analyzed for each experimental group). G) Quantification of BFU-E colonies in mouse fetal liver erythroid precursor cells from Samd14 conditional knockout mouse that express recombinant full length Samd14 (full length), S14ΔSAM (missing SAM domain), S14R381Q (PBM mutant) or an empty vector (EV). Error bars represent standard error of mean (SEM). *p<0.05; **p<0.01; ****p<0.0001 (two-tailed unpaired Student’s t test). H) Molecular mechanism promoting BFU-E activity involves a Samd14-PI3P interaction via the Samd14 SAM.

### Samd14 and PI3P coordinate erythroid differentiation after hemolytic anemia

PI3P is a relatively low-abundant lipid moiety in cells localized to early endosomes and in autophagic membranes (Wallroth & Haucke, 2018). Our discovery that Samd14 binds specifically to PI3P and not PI(3,5)P_2_ in the PIP strip assay hints at Samd14 localization to early endosomes and eventual release as they transition into late endosomes. If Samd14 localized to endosome/autophagosome membranes to regulate autophagy, we reasoned that autophagy induced by serum starvation may relocate Samd14 to associate with intracellular organelles. To test this, we performed subcellular fractionation of crude organelles after serum starvation in human erythroleukemia (HEL) cells (**Figure 6A**). We detected a relative increase in the amount of Samd14 present in the organelle compartment 4 hours post serum-starve (**Figure 6B)**. RAB5, LC3, CAPZB and GAPDH were used to validate separation of the organelle fraction (**Figure 6C**). To test whether this phenomenon depends on PI3P synthesis, which is catalyzed by the VPS34 enzyme, we treated cells with the VPS34 inhibitor SAR405 (Ronan *et al*, 2014). Inhibiting PI3P synthesis prevented Samd14 localization to organelles (**Figure 6D-6E**).

**Figure 6.**
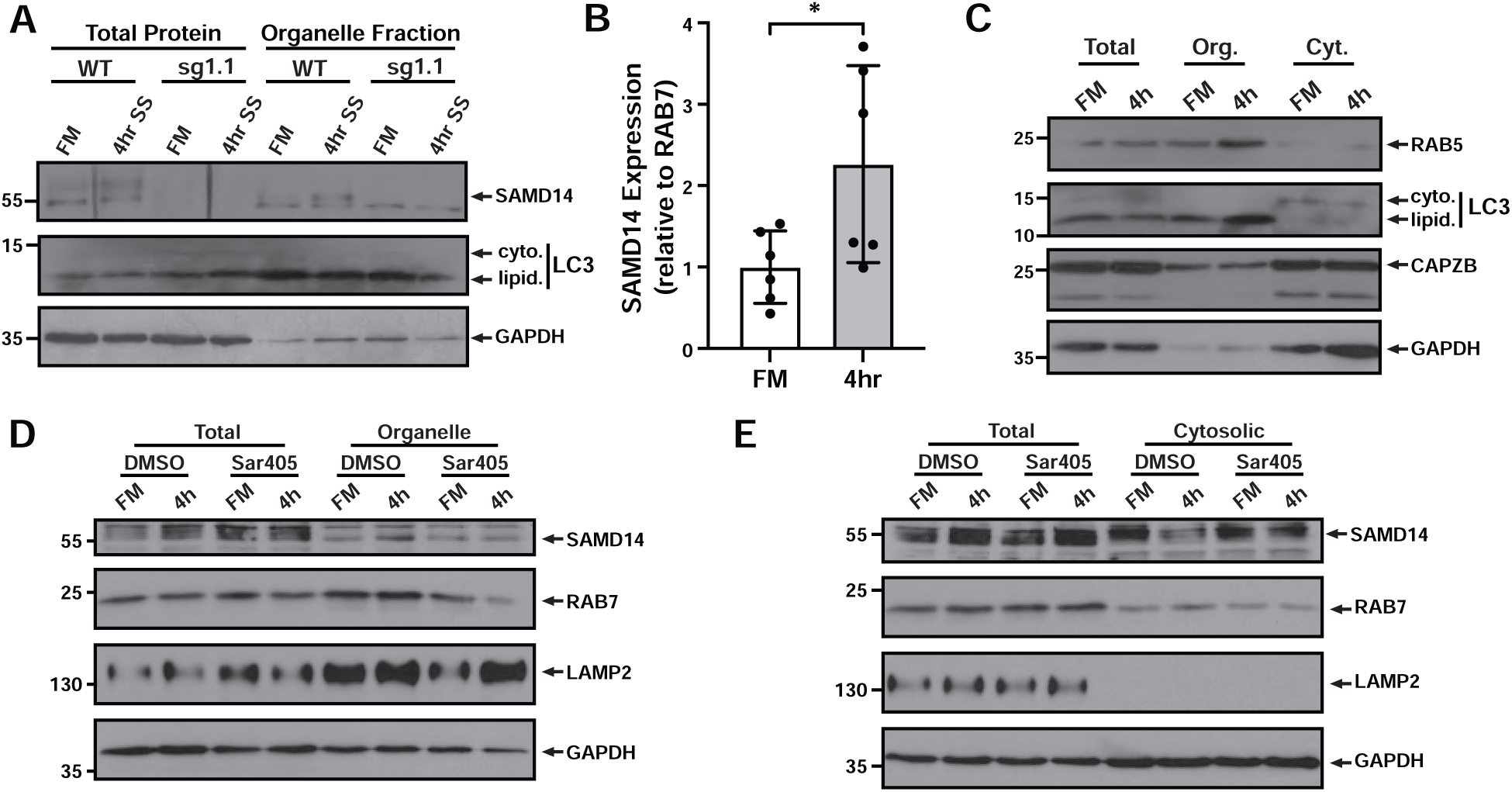
Samd14 localizes to organelles in starvation-induced autophagy. A) Western blotting of total and fractionated protein lysates from control or Samd14-deficient human erythroleukemia (HEL) cells after 4 hours serum starvation (SS) or full media (FM) controls. B) Semi-quantitative densitometry of SAMD14 protein levels in organelle fractions comparing FM vs. SS conditions (N=6). Error bar represents standard deviation); *p<0.05. C) Western blotting of protein lysates from total, organelle (Org.) and cytoplasmic (Cyt.) fractions in FM and SS-treated cells. D) Western blotting of protein lysates from total and organelle fractions in FM and SS-treated cells pre-incubated with Sar405 (1uM). E) Western blotting of protein lysates from total and cytosolic fractions in FM and SS-treated cells pre-incubated with Sar405 (1 µM).

To test the role of PI3P and VPS34 activity in erythropoiesis, primary erythroid precursors derived from WT mouse embryonic day 14.5 (E14.5) fetal liver were treated with SAR405 for 72 hours (**Figure 7A**). SAR405 treated fetal liver cells grew at a slower rate (**Figure 7B**) and cell pellets were paler in color compared to DMSO-treated controls (**Figure 7C**). Cell surface markers CD71 and Ter119 were used to track erythropoiesis over 3 days in culture (Koulnis *et al*, 2011) (**Figure 7D**). VPS34 inhibition decreased the percentage of committed erythroid cells (CD71^high^TER119^-^) by 1.5-fold (p=0.001), dose 5 µM) commensurate with increased percentages of maturing erythroid cells (CD71^+^TER119^+^) by 1.3-fold (p=0.05) (**Figure 7E**). Strikingly, mature erythroid cells (CD71^low/neg^TER119^+^) decreased by 2.2-fold (p=0.01) (**Figure 7E**). R3 erythroblasts of SAR405 treated groups contained a higher fraction of Kit positive cells (5.1-fold, p=0.01) (**Figure 7F**). SAR405-treated cells expressed 2.5-fold more *Gata2* mRNA, and 2.3-fold and 1.9-fold lower globin (*Hba-a2*) and Glycophorin A (*Gypa*) mRNA compared to controls, respectively, consistent with a less differentiated culture (**Figure 7G**). Taken together, treatment of differentiating erythroblasts with a selective VPS34 inhibitor impairs terminal erythropoiesis.

**Figure 7.**
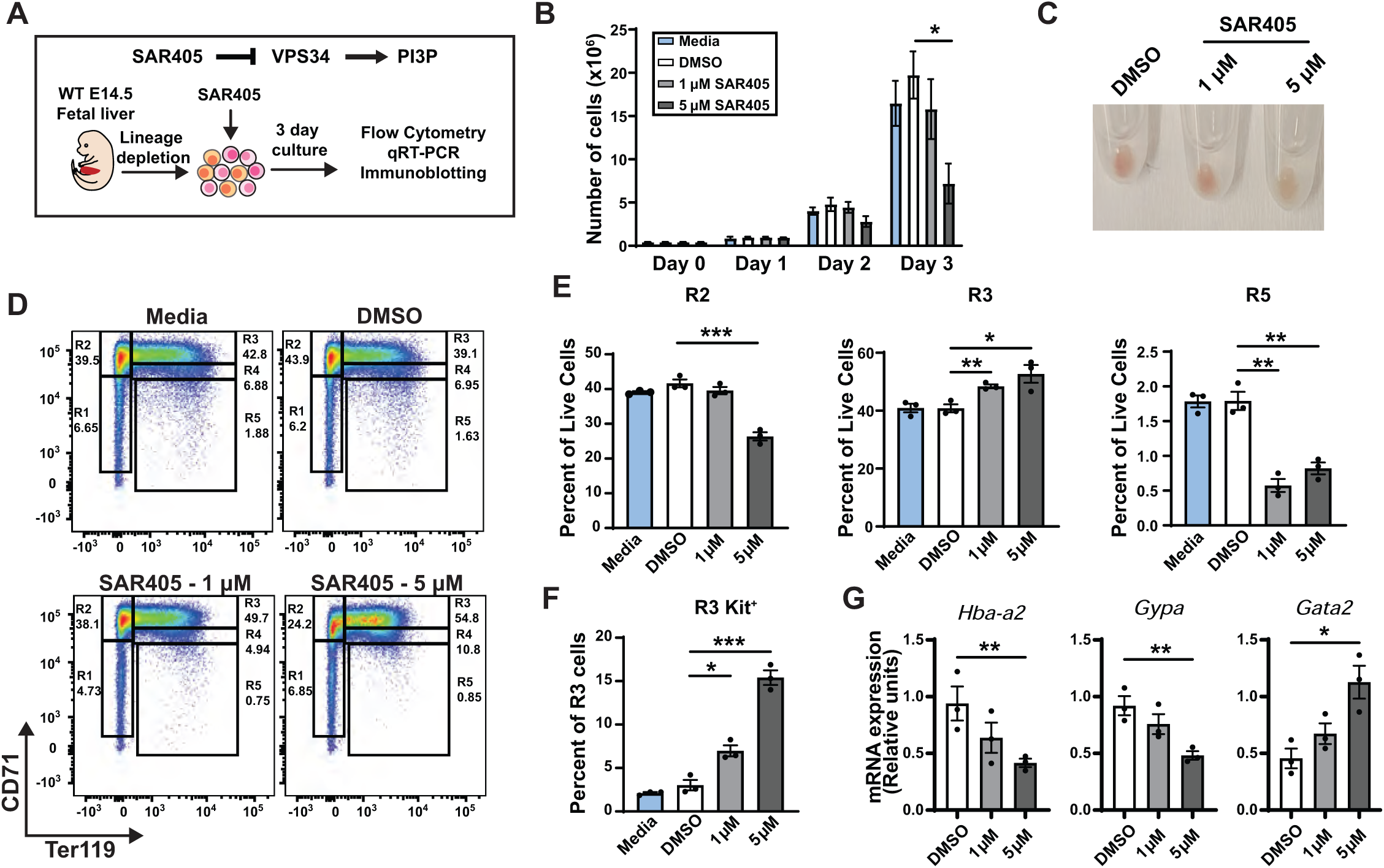
VPS34 promotes terminal erythropoiesis in E14.5 mouse fetal liver erythroid precursor cells. A) Experimental layout. Cells were isolated from WT E14.5 mouse fetal liver treated with DMSO or SAR405 (1 µM and 5 µM). B) Number of WT primary cells in expansion culture over a 3-day time course treated with indicated doses of SAR405 (or DMSO vehicle control) determined by trypan-negative cells (N=3). C) Cell pellets from SAR405-treated cultures (day 3). D) Representative flow cytometry plots of cells stained with erythroid markers CD71 and TER119. Populations R1-R5 represent erythroid cells in progressive stages of maturation. E) Quantitation of flow cytometry data in (D). N=5 embryos per group. F) Quantitation of the percentage of CD71^+^Ter119^+^ (R3) cells that were Kit^+^. G) Quantitation of *Hba-a2*, *Gypa*, and *Gata2* mRNA levels in cells treated with 1 or 5 µM SAR405 or DMSO controls for 3 days (N=3). Error bars represent standard error of mean (SEM). *p<0.05; **p<0.01; ***p<0.001 (two-tailed unpaired Student’s t test).

Lastly, we asked whether there were unique requirements for PI3P in the context of stress erythropoiesis. WT and *Samd14*-KO mice were treated with PHZ for three days prior to isolation and *ex vivo* culture of splenic erythroid precursors (**Figure 8A**). SAR405 treated cells were paler in color compared to DMSO-treated controls (**Figure 8B**). SAR405 treated cells contained a smaller percentage of CD71^+^Ter119^+^ cells than control populations, and an accumulation of less differentiated CD71^+^Ter119^-^ cells (**Figure 8C**). In Samd14-deficient cells, the percentage of CD71+Ter119+ cells after SAR405 treatment were consistently higher at both doses, indicating that Samd14 expression sensitizes erythroid precursors to a SAR405-induced block in erythropoiesis (**Figure 8D**). In the context of SAR405 treatment, Samd14-deficient cells have less lipidated LC3-II relative to WT cells (**Figure 8E**). This demonstrates that SAR405 robustly inhibits autophagy in WT erythroid precursor cells, and this is attenuated in Samd14-deficient cells (**Figure 8F**). Samd14 interacts with PI3P, a crucial lipid in cellular autophagy, to modulate autophagy and erythropoiesis in the context of acute anemia stress.

**Figure 8.**
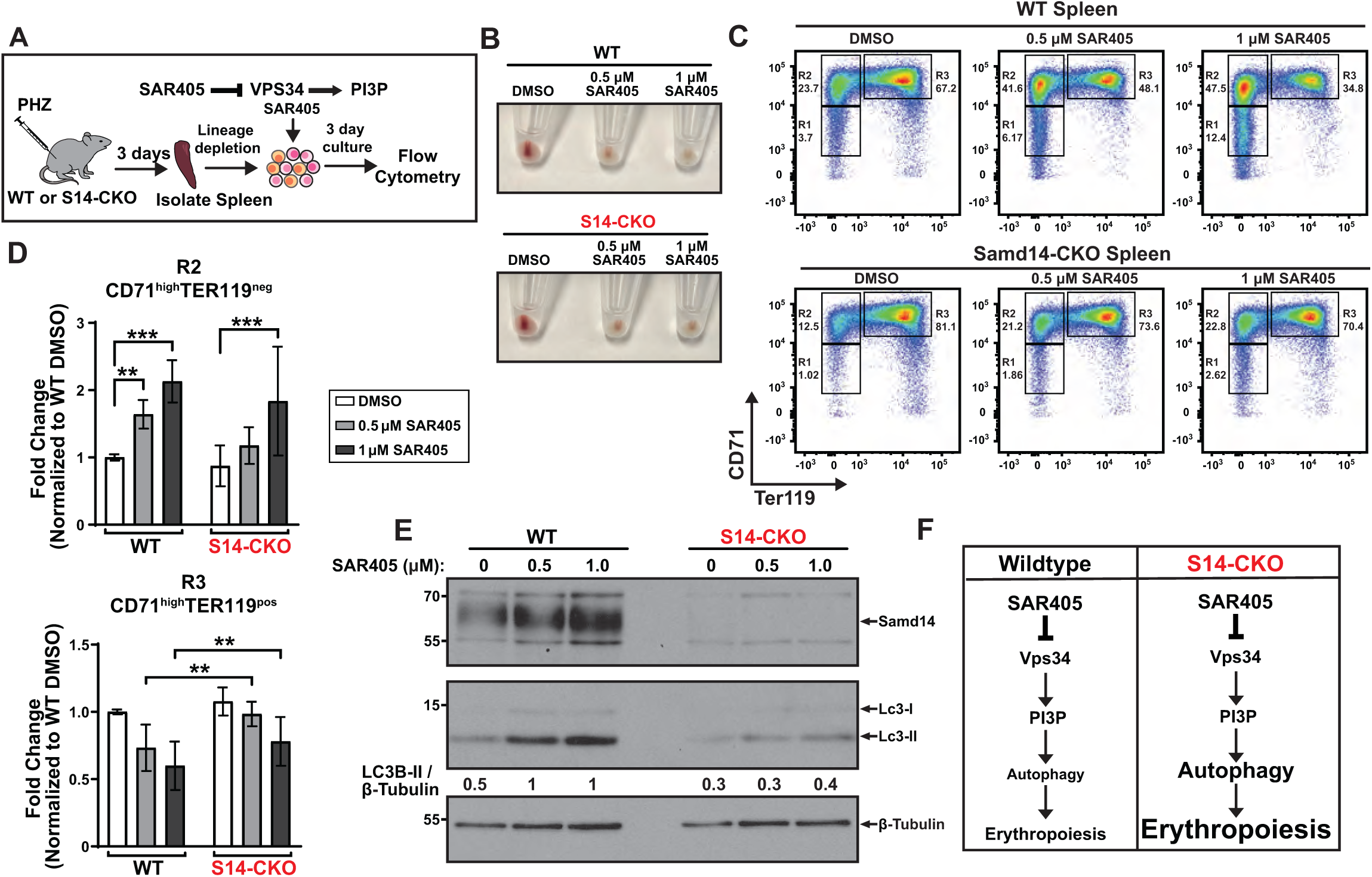
SAMD14 expression sensitizes splenic erythroid precursor cells to VPS34 inhibition. A) Experimental layout of splenic EPCs isolated from Samd14^fl/fl^ (WT) and *Samd14*^fl/fl;Vav-Cre^ (S14-CKO) mice treated with VPS34 inhibitor SAR405 for 3 days post PHZ-induced anemia. B) Cell pellets from SAR405-treated cultures (day 3). C) Representative flow cytometry plots showing progressive stages of maturing erythroid cells in presence and absence of SAMD14 and SAR405. D) Quantitation of flow cytometry data in (C) for R2 (CD71^high^TER119^neg^) and R3 (CD71^high^TER119^pos^) populations. ‘Percentage of live’ was expressed as a fold change over average value for WT DMSO treated cultures (a total of n = 7 mice per group). Error bars represent standard deviation (SD). **p<0.01; ***p<0.001; ****p<0.0001 (Two-way ANOVA). E) Western blotting protein lysates from WT and S14-CKO cells after 3 days culture n DMSO or SAR405-treated cultures. F) A current working model depicting the Samd14-dependent sensitization of erythroid cells to VPS34 inhibition.

## Discussion

The identification of a lipid interacting partner for the Samd14-SAM domain has resolved a puzzling gap in knowledge regarding Samd14 molecular mechanisms to restore homeostasis in anemia. The PI3P lipid moiety serves as a docking site for Samd14 and other proteins which are capable of both promoting and opposing the tightly regulated process of autophagy (Obara & Ohsumi, 2008). Prior reports have shown that PI3P interacts with another Samd14-interacting protein partner, the actin CP complex, to control actin polymerization (Mi *et al*., 2015). Our data shows that Samd14 interacts with both PI3P and all three subunits of the CP complex (Capzα1, Capzα2, Capzβ) (Ray *et al*., 2022), forming a multi-subunit complex at endosome/autophagosome membranes. As PI3P is a spatial coordinator of signaling molecules and proteins involved in membrane formation, and removal of Samd14 alters autophagic responses, this suggests that the PI3P-Samd14-CP complex plays a role in autophagy.

Autophagy is closely coupled with cell signaling pathways controlled by Samd14. As a pro-proliferation and survival pathway, the SCF/Kit receptor tyrosine kinase (which is facilitated by Samd14 expression) activates PI3K and AKT signaling, which generally leads to less autophagy in cells (Ames *et al*, 2023; Engelman *et al*, 2006; Ertmer *et al*., 2007). In other contexts, including Kit-expressing gastrointestinal stromal tumors, Kit signaling suppresses autophagy (Hsueh *et al*, 2014). Oncogenic Kit mutations have also been shown to induce autophagy in leukemic cells (Larrue *et al*, 2019). While direct evidence for PI3P-binding effector proteins (other than Samd14) in Kit signaling is lacking in the literature, PI3P can control signaling through epidermal growth factor receptor (EGFR) (Petiot *et al*, 2003). PI3P influences the duration of downstream signaling through the MAPK pathway (Taub *et al*, 2007). Our phospho-proteomics data, and other evidence linking the Kit signaling facilitator Samd14 and its interacting protein/lipid to autophagy, support the idea that SCF/Kit signaling suppresses autophagy in erythropoietic contexts. Autophagy responses to pro-recovery and pro-differentiation signaling pathways are essential for maintaining programs of terminal differentiation in erythropoiesis, and in stress contexts.

Cells are capable of initiating autophagy responses during stress without causing cell death. Pro- and anti-autophagy mechanisms are needed to control this balance (Cecconi & Levine, 2008). Both excessive and ineffective autophagy can cause anemia (Zhang *et al*., 2015). Samd14 is one of several autophagy regulators previously linked to erythropoiesis in normal and stress contexts, in which targeted deletion in mice yields anemia phenotypes. Other examples include the ATG4 protein, which controls autophagosome fusion in late, differentiating erythroblasts (Betin *et al*, 2013), ULK1 which is needed for the effective clearance of mitochondria in reticulocytes (Kundu *et al*, 2008), and FIP200 which is an autophagy-promoting protein that is needed during embryonic development for selective autophagy (Liu *et al*, 2010). Transcriptional control of autophagy has been well established to occur downstream of the GATA1 transcription factor (Kang *et al*., 2012; Welch *et al*, 2004). Autophagy-related proteins were upregulated in erythroid progenitors after acute anemia, several of which were downregulated at the RNA level, suggesting potential feedback loops. The distinct requirements for Samd14 expression in the context of red blood cell regeneration led us to propose a model in which GATA factor-mediated Samd14 upregulation acts as a brake on excessive autophagy in acute anemia, thereby promoting cell and organismal survival. Our findings add to a body of knowledge surrounding autophagic control in erythropoiesis, implicating the GATA factor-regulated Samd14 protein as a relevant modulator of autophagy during erythropoiesis.

## Materials and Methods

### Plasmid Constructs and Site Directed Mutagenesis

HA-tagged full-length Samd14, Samd14 lacking the SAM domain (S14ΔSAM), Samd14 lacking the capping protein binding (CPB) domain (Samd14 ΔCPB), and Samd14 lacking both SAM and CPB domains (S14ΔCS) (Ray *et al*., 2022) were cloned into mammalian expression plasmid pMSCV-Puro-IRES-GFP (Addgene #21654) using BglII and EcoRI restriction digestion sites. To generate Samd14 lacking the polybasic basic motif (S14R381Q), the full length Samd14 construct was mutated at arginine position 381 to glutamine using Phusion Hot Start II High Fidelity PCR Master Mix. Details regarding site directed mutagenesis is available upon request. Flag-tagged full length Samd14 cloned into pMSCV-Puro-IRES-mCherry was purchased from Vector Builder. All retroviral MSCV plasmids were packaged with pCL-Eco (Addgene #12371) in 293T cells using calcium phosphate transfection.

### Mice

Samd14-Enh knockout (S14^ΔE/ΔE^) mice were generated by site-directed TALEN deletion (Hewitt *et al*., 2017). Samd14 conditional knockout (S14-CKO) mice was generated by crossing Samd14 floxed mice (Schaefer *et al*., 2023) to the hematopoietic-specific Vav-Cre line (Jackson Lab strain 008610). All animal experiments met the ethical standards set by the Institutional Animal Care and Use Committee of the University of Nebraska Medical Center.

### Phospho-proteomics

WT and (S14^ΔE/ΔE^) spleens were manually dissociated 3 days post-PHZ injection. Kit+ cells were isolated with biotin-conjugated anti-mouse CD117 (Kit) antibody and MojoSort streptavidin-conjugated magnetic nanobeads (Biolegend). 10^7^ Kit^+^ cells per condition (N=3 mice per genotype) were serum-starved for 2 hr in 1% BSA/IMDM at 37°C and subsequently treated with 10 ng/ml SCF or vehicle (PBS) for 25 mins. 100 μg of protein was alkylated by addition of 8 μl 0.5M IAM in water for 40 min and acetone precipitated overnight at -20°C. Trypsin digested protein lysates were checked for extent of digestion >95% and TMT labeled according to the TMTpro Mass Tag Labeling Reagents and Kits User Guide. Labeled peptides were phosphopeptide enriched on titanium beads. Elution from the titanium beads was carried out with 5% ammonium hydroxide and 5% pyrrolidine. Phosphopeptide fractions were loaded onto a 75 μm x 2 cm Acclaim PepMap 100 C18 trapping column in 1.5% acetonitrile, 0.05% TFA at a flow rate of 5 μl/min using a RSLC nano Ultimate 3000 chromatographic system (Thermo Scientific). The samples were run twice using 2 different sets of 3 compensation voltages: -40, -60, -80 and -50, -70, -90. FAIMS-CID-MSA MS2 (in the ion trap MS) with RTS SPS MS3 (in the Orbitrap MS). Raw mass spectrometry data was analyzed using Proteome Discoverer 2.4 (ThermoFisher). Identified phospho-sites and abundances were compared using RokaiXplorer against the reference mouse proteome.

### Primary Cell Culture

To induce hemolytic anemia, 8- to 12-week old mice were subcutaneously injected with a single dose (100 mg/kg) of phenylhydrazine (Sigma Aldrich) freshly resuspended in sterile phosphate buffered saline (PBS). After 72 hours, spleens harvested from WT (*Samd14^fl/fl^*), S14-CKO mice or S14^ΔE/ΔE^ mice were dissociated in PBS supplemented with 2% fetal bovine serum (FBS). Single-cell suspensions were obtained by passing cells through a 35 μm nylon filter. Cells were labeled with biotin-conjugated antibodies against lineage markers of T-cells (anti-mouse CD34ε; clone 145–2 C11), macrophages (anti-mouse CD11b; clone M1/70), B-cells (anti-mouse CD19; clone 6D5 and anti-mouse CD45R/B220; clone RA3-6B2), granulocytes (anti-mouse Gr-1; clone RB6-C5) and erythrocytes (anti-mouse Ter119). Erythroid precursors were enriched by negative selection of biotinylated cells using MojoSort streptavidin-conjugated magnetic nanobeads (Biolegend). The resultant lineage depleted cells were cultured in erythroid expansion medium: StemPro-34 (Invitrogen) containing 2 mM L-glutamine, 1% Penicillin-Streptomycin, 0.1 mM monothioglycerol, 1 µM dexamethasone, 0.5 U/mL erythropoietin, and 1% mSCF Chinese Hamster Ovary cell conditioned medium. For retroviral infections, 100 μL viral supernatant was added to 1×10^6^ lineage depleted cells along with polybrene (8 μg/ml) and HEPES buffer (10 μl/ml) in a 12-well plate. Cells were spinoculated at 2600 rpm for 90 min at 30 °C and thereafter maintained at a density of 0.5×10^6^ cells/mL for up to 72 hours.

### G1E Cell Culture

Mouse embryonic stem cell-derived *Gata1*-null Erythroid (G1E) cells (gift from the Mitch Weiss lab) resemble normal proerythroblasts (Weiss *et al*, 1997). Cells were maintained in Iscove’s DMEM supplemented with 15% FBS, 0.5% mSCF-conditioned media from Chinese Hamster Ovary cells, 0.1 mM monothioglycerol (Sigma) and 2 U/mL erythropoietin (Amgen). The Samd14 intronic enhancer was knocked out in cells (G1E-ΔEnh) by TALEN-mediated deletion. Full length or mutant Samd14 expression was restored by retroviral spinoculation (2600 rpm for 90 min at 30 °C) of 1×10^6^ G1E-ΔEnh cells with 100 μL viral supernatant, polybrene (8 μg/ml) and HEPES buffer (10 μg/ml). Cells were maintained at 0.5×10^6^ cells/mL density for 48 hours after which FACS-purification of GFP^+^ cells was performed on a FACSAria (BD Life Sciences).

### SAM Dimerization Assay

For SAM-SAM dimerization assay, we generated G1E-ΔEnh cells expressing both HA-tagged and Flag-tagged full length Samd14. A second experimental group comprised cells expressing HA-tagged SAM-deleted Samd14 together with Flag-tagged full length Samd14. Negative controls included cells expressing either HA-tagged Samd14 or Flag-tagged Samd14 or the empty vector. Either GFP or mCherry or double positive cells were sorted in a FACSAria and expanded for up to a week. For HA pulldown experiments, 3 × 10^7^ sorted cells expressing different Samd14 constructs were harvested and lysed for an hour in 1% or 0.1% NP-40 lysis buffer (150 nM NaCl, 50 mM Tris pH 8.0, 2 mM DTT, 0.2 mM PMSF, 20 µg/ml leupeptin) at 4 °C. Lysates were incubated with HA-agarose beads (Pierce) overnight, washed and boiled with SDS lysis buffer. Recombinant protein expression was detected by western blotting using anti-rabbit HA (Cell Signaling Technology; 1:2000) and anti-rabbit DYKDDDDK tag antibody (Cell Signaling Technology; 1:4000).

### PIP Strip

HA-tagged recombinant proteins were immunoprecipitated with HA-agarose beads (Pierce). Once HA-tagged proteins were eluted from the bead with synthetic HA peptide (Pierce #26184), samples were passed through Vivaspin500 10kDa MWCO columns (Cytiva #28932225) to remove 1.1 kDa HA peptide while retaining the 50-55 kDa SAMD14 protein. To control for correct completion of steps involved in this assay, we spotted approximately 150 ng protein and 0.5 uL of primary and secondary antibody directly onto open areas of the dry membrane. Spots were allowed to dry completely before blocking the membrane with 1% non-fat dry milk in TBST. Membranes were incubated with 2 µg/mL purified protein in 1% milk/TBST followed by incubation with anti-rabbit SAMD14 primary antibody and anti-rabbit HRP conjugated secondary antibody. The blots were probed with ECL substrate and developed using X-ray films.

### Proximity Ligation Assay (PLA) and Confocal Microscopy

PLA was performed using the Duolink PLA kit (Sigma Aldrich). FACS purified GFP^+^ G1E cells expressing recombinant SAMD14 constructs were cytospun onto charged slides at 800g for 5 minutes at low acceleration such that each spot had 50,000 cells. The slides were fixed, permeabilized and stained according to the manufacturer’s instructions. We used primary antibodies against mouse PI3P (Echelon Biosciences; 1:300) and rabbit HA-tag (Cell Signaling Technology; 1:1200). Images were acquired by confocal laser scanning microscopy (Zeiss 880) and PLA puncta was quantified using the ImageJ plugin Andy’s Algorithm (Law *et al*, 2017).

### Colony formation assay

Fetal liver isolated from day 14.5 embryos were dissociated in PBS containing 2% FBS and passed through a 35 μm nylon filter to obtain single-cell suspensions. Post lineage depletion, fetal liver erythroblasts were retrovirally infected to express either empty vector (EV), Samd14, S14ΔSAM or S14R381Q constructs. After 40-42 hours of culture in erythroid expansion media, GFP+ cells were purified using FACS. Sorted GFP^+^ cells were mixed with Methocult M3434 (STEMCELL Technologies) containing Epo, SCF, IL-3 and IL-6 and plated in 3 replicate wells of a 12-well plate (5 × 10^3^ cells in 0.5 mL). BFU-E colonies were counted 6 days after culturing.

### Subcellular fractionation

HEL 92.1.7 cells (ATCC) were grown in full media (RPMI 1640, 10% FBS, 1% Pen/Strep) for 24 hours. Cells were starved in RPMI 1640 for 4 hours prior to fractionation. SAR405 treated cells (Selleck Chemicals #S7682) were grown for 24 hours in full media supplemented at 1µM or an equal volume of DMSO. This was followed by 4-hour starvation in RPMI 1640 alone containing SAR405 or DMSO. Cells were harvested using the Minute™ Plasma Membrane/Protein Isolation and Cell Fractionation Kit (Invent Biotechnologies, Plymouth, MN). Sample fractions were lysed for 30 minutes on ice in a mild lysis buffer (50mM Tris-HCl, 150mM NaCl, 1mM EDTA, 1% Triton-X, pH 7.5) in presence of protease inhibitor cocktail (Roche cOmplete™, Mini, EDTA-free Protease Inhibitor Cocktail). Samples were centrifuged at max speed for 10 minutes at 4°C and supernatant was collected for BCA protein quantification assay (Pierce #23227). Equal amounts of protein were loaded in an SDS-PAGE and analyzed by western blotting.

### Flow cytometry

Mouse primary cells were stained with fluorophore-conjugated antibodies (all from BioLegend) specific for surface proteins CD71 (R17217), Ter119 (116212), Kit (clone 2B8) (105814) for 45 minutes to an hour on ice. Cells were washed once with PBS, resuspended with the viability dye DAPI and analyzed with BD LSRII Y/G or BD LSRFortessa (BD Life Sciences). Data analysis was performed with FlowJo 10.10.0.

### Western Blotting

Protein lysates were prepared by boiling in sodium dodecyl sulfate (SDS) lysis buffer (25mM Tris pH 6.8, 2% b-mercaptoethanol, 3% SDS, 5% bromophenol blue, and 5% glycerol) for 10 minutes at 95 °C. Proteins were resolved by SDS-PAGE and detected with Pierce ECL Western blotting substrate (Thermo Scientific) using X-ray film processor. Primary antibodies used are as follows: polyclonal rabbit anti-Samd14 (Hewitt lab), rabbit anti-HA (Cell Signaling Technology), anti-β-Actin (Cell Signaling Technology), anti-LC3A/B (Cell Signaling Technology), anti-SQSTM1/P62 (Cell Signaling Technology), anti-β-Tubulin (Cell Signaling Technology), anti-phospho-AMPKα (Thr172) (Cell Signaling Technology), anti-AMPKα (Cell Signaling Technology), anti-phospho-eIF2α (Ser51) (Cell Signaling Technology), anti-eIF2α (Cell Signaling Technology), anti-phospho-4E-BP1 (Thr37/46) (Cell Signaling Technology), anti-HA-Tag (Cell Signaling Technology), anti-Flag DYKDDDDK Tag (Cell Signaling Technology), anti-CapZ-β (Santa Cruz), anti-GAPDH (Santa Cruz), anti-Rab5 (Santa Cruz), anti-Rab7 (Cell Signaling Technology), anti-LAMP2 (Santa Cruz). Secondary antibodies include goat-anti-mouse-IgG-HRP (Jackson Labs) and goat-anti-rabbit-IgG-HRP (Jackson Labs).

### Quantitative PCR

RNA was extracted from 1 to 2 × 10^6^ cells with 1 mL TRIzol (ThermoFisher). For cDNA synthesis, 1 µg of RNA was mixed with 200 ng oligo(dT) primers and 50 ng random hexamer primers and incubated at 68 °C for 10 minutes. Reverse transcription was carried out using M-MLV reverse transcriptase (NEB) in the presence of 10 mM DTT, RNasin (Promega), and 0.5 mM dNTPs at 42 °C for 1 hour, followed by heat inactivation at 98 °C for 5 minutes. Quantitative real-time PCR (qPCR) was performed on a QuantStudio 3 system (Applied Biosystems) using 2 µl of cDNA, Power SYBR Green Master Mix (ThermoFisher), and 200 nM gene-specific primers on the Quant Studio 3 real time PCR system (Applied Biosystems). Relative expression levels were determined from a standard curve generated with serial dilutions of control cDNA. Control reactions lacking reverse transcriptase were included to confirm the absence of genomic DNA contamination.

## Acknowledgments

We are grateful to the core facilities at UNMC and the support these facilities receive from UNMC. Notably, we extensively used the UNMC Flow Cytometry Research Facility for this project, which is supported by the Nebraska Research Initiative (NRI) and The Fred and Pamela Buffett Cancer Center’s (FPBCC) National Cancer Institute Cancer Support Grant, and the Advanced Microscopy core facility. This study was supported by National Institutes of Health/National Heart, Lung, and Blood Institute [R01 HL155439] and Nebraska Stem Cell Research (DHHS 2019-01). Phospho-proteomics was conducted in consultation with Dr. Sophie Alvarez and Mike Naldrett in The Proteomics & Metabolomics Facility (RRID:SCR_021314), Nebraska Center for Biotechnology at the University of Nebraska-Lincoln, supported in party by the Nebraska Research Initiative. We declare no competing financialinterests.

